# Mechanisms of force transmission on cytokinesis Formin Cdc12 in fission yeast revealed by coiled-coil force sensors

**DOI:** 10.1101/2025.05.14.653946

**Authors:** Takumi Saito, Yuan Ren, Julien Berro

## Abstract

Cytokinesis is a fundamental process in cell division, where an actomyosin contractile ring plays a central role in completing the cell division. Although some experimental and computational efforts have evaluated ring tension and the molecular organization of rings, the mechanisms of force transmission at the molecular level remain unclear. Here, we used our novel coiled-coil force sensors to measure the force distribution along the formin Cdc12, a key cytokinesis protein in fission yeast. Our force measurements revealed that individual formin Cdc12 molecules transmit up to ∼6 pN with distinct mechanisms between the regions upstream and downstream of the formin homology 2 (FH2) domain, which binds the barbed ends of actin filaments. The force transmitted on the N-terminal region before the FH2 domain requires anchoring to a cytokinetic node via the Cdc15 binding region, but is independent of the C-terminal region after FH2. In contrast, the region after the FH2 domain transmits forces independently of the Cdc12’s N-terminal region. Force screening using the coiled-coil force sensors found that a region in Cdc12’s disordered C-terminal tail can associate with the contractile ring. Altogether, the force measurements with our coiled-coil force sensors allowed us to uncover the mechanisms of force transmission along Cdc12 and characterize new motifs and binding partners.

## Introduction

Cytokinesis is a fundamental process of cell division in animals, fungi, and amoebas, where an actomyosin contractile ring assembles and constricts to pinch the mother cells into two daughter cells (Pollard and O’Shaughnessy, 2019). Numerous efforts have been identifying cytokinesis proteins and unveiling their molecular organizations in contractile rings (Mangione and Gould, 2019). In particular, myosin motors produce contractile forces by pulling actin filaments that are nucleated by Cdc12 formins, which are anchored to the membrane via structures called nodes. However, the mechanisms of force transmission from the actomyosin ring to the plasma membrane remain unclear. Understanding which proteins of the ring bear forces and measuring the magnitude of these forces will allow to better understand how actomyosin rings produce and transmit forces.

The fission yeast *Schizosaccharomyces pombe* is a model organism often used for studying cytokinesis due to well-characterized and well-conserved cytokinesis molecules and processes. In fission yeast, cytokinetic nodes containing myosin-II Myo2p, myosin light chains Rlc1 and Cdc4, IQGAP Rng2, F-BAR protein Cdc15, and formin Cdc12, among other proteins, are assembled in a broad band on the plasma membrane in the middle of the cell (Wu *et al*., 2003; Coffman *et al*., 2009; Laporte *et al*., 2011; Lee *et al*., 2012; Laplante *et al*., 2016). The actin filaments nucleated by Cdc12 molecules of one node are captured by myosin-II motors from other nodes to align them into a thin ring using a search-capture-pull-release (SPCR) mechanism, over ∼10 min (Vavylonis *et al*., 2008; Lee *et al*., 2012). During anaphase B, the ring matures for ∼25 min while keeping a constant diameter before it constricts (Wu *et al*., 2003). During anaphase, a ring tension of ∼390 pN was measured in fission yeast protoplasts, yeast cells lacking a cell wall (Stachowiak *et al*., 2014). As constriction progressed, the ring tension increased to ∼800 pN with a mean of 640 pN (Mcdargh *et al*., 2021). The ring tension is produced by the interactions between actin and three myosins, the myosin-IIs Myo2 and Myp2, and the myosin-V Myo51. Myo2 join the nodes at their inception during the broad band formation, whereas Myp2 and Mo51 are recruited to the ring during the maturation phase (Laplante *et al*., 2015). Although the molecular organization and the average ring tension has been measured, how forces can be transmitted along the ring is still unclear, especially considering the unique molecular organization in contractile rings, which is different from sarcomeric structures in skeletal muscles and in stress fibers of non-muscle cells,.

Fission yeast cytokinetic formin Cdc12, an actin polymerization protein, is essential for cytokinesis, are present from the beginning of node assembly to the end of ring constriction (Wu and Pollard, 2005; Yonetani *et al*., 2008; Pollard and O’Shaughnessy, 2019; Homa *et al*., 2021) (Fig. 1A). Cdc12 promotes actin filament elongation, via the two conserved formin homology (FH) domains (Breitsprecher and Goode, 2013). The polyproline-rich FH1 domain captures and delivers profilin-bound actin monomers to actin filaments via the FH2 domain (Otomo *et al*., 2005; Courtemanche and Pollard, 2012; Courtemanche *et al*., 2013; Homa *et al*., 2021). FH2 domains form a donut-shaped homodimer that caps the barbed end of an actin filament, to which incoming actin monomers can be added (Kovar *et al*., 2003; Paul and Pollard, 2009). In addition to the actin-binding domains, the N-terminal region of Cdc12 interacts with the F-BAR protein Cdc15p, which is another component of cytokinetic nodes at the plasma membrane (Carnahan and Gould, 2003; Willet *et al*., 2015; Snider *et al*., 2020). Considering the domain organization of formins, we hypothesized that Cdc12 directly transmits the forces generated by myosins pulling on FH2-bound actin filaments to the nodes via Cdc15 (Fig. 1B). Other non-mutually exclusive hypotheses are that force could be transmitted to the nodes or to other components of the contractile ring via other domains of Cdc12. To test these hypotheses, we used our recently developed coiled-coil force sensors to detect and measure the forces along Cdc12 (Ren *et al*., 2025b).

**Figure 1.**
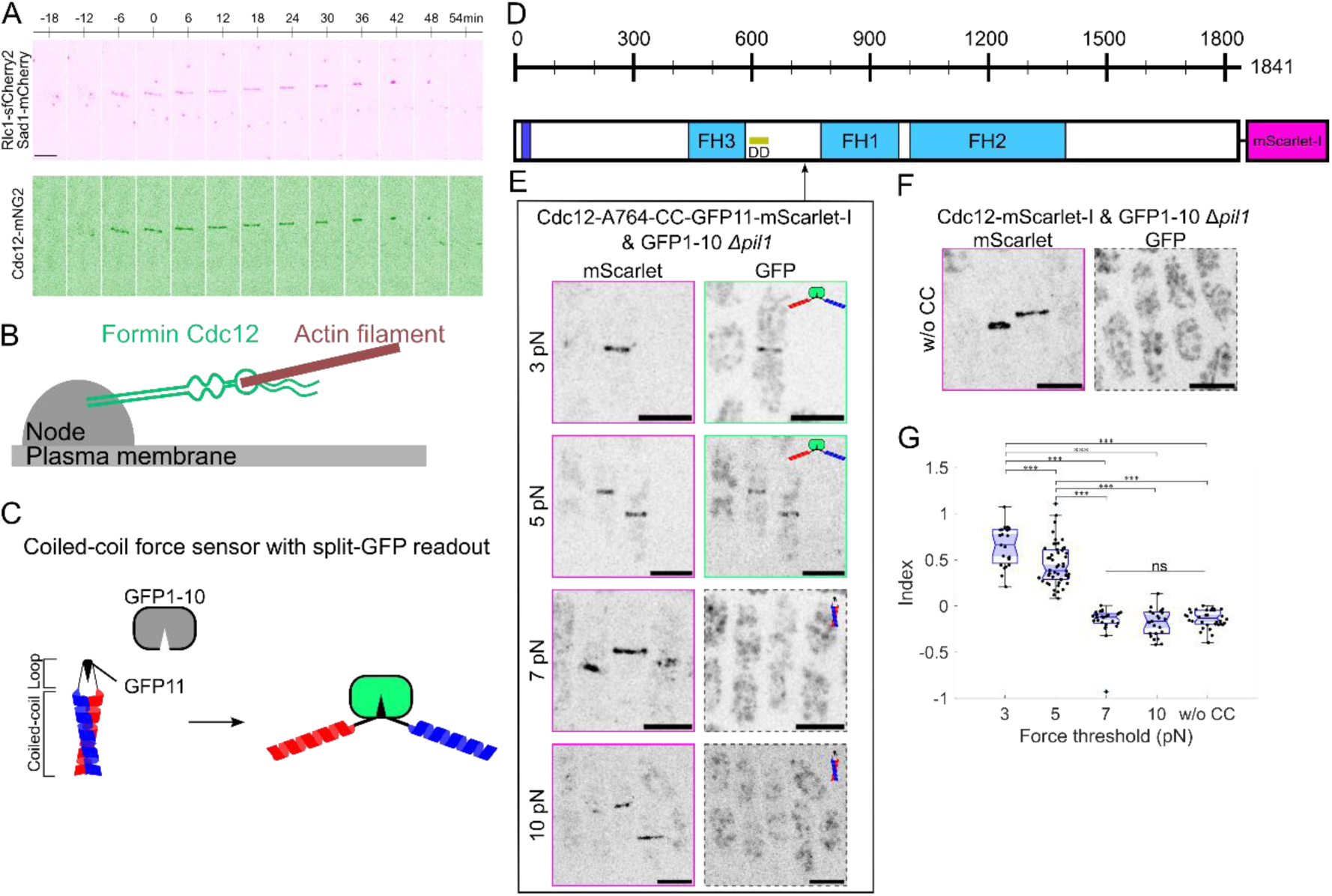
Force measurements on the formin Cdc12 during cytokinesis in fission yeast *S. pombe*, using the coiled-coil (CC) force sensors. **A** Time lapse images of the fluorescently tagged formin Cdc12 (Cdc12-mNeonGreen2), Rlc1 (Rlc1-sfCherry2) as a ring marker, and Sad1 (Sad1-mCherry) as a spindle pole body marker. Scale bar; 10 µm. **B** The formin Cdc12 is anchored to a node on the plasma membrane and binds the barbed end of actin filaments. **C** Schematics of the coiled-coil force sensors with the split-GFP readout. **D** The domain organization of the fluorescently tagged formin Cdc12 (Cdc12-mScarlet-I) composed of the N-terminal node-binding region, the formin homology (FH) domains, and the putative dimerization domain (DD). **E** Fluorescence images of fission yeast expressing fluorescently tagged Cdc12 (Cdc12-mScarlet-I) and GFP1-10 replacing *pil1* gene. The 3, 5, 7, or 10 pN force sensors with the GFP11 linker are genetically inserted at A764 of Cdc12-mSCarlet-I. Scale bars; 10 µm. **F** Fluorescence images of the control strain expressing Cdc12-mScarlet-I and GFP1-10 replacing *pil1* gene. Scale bars; 10 µm. **G** Quantification of the force measurements by the force index, represented as mean ± SD (*n*=23, 52, 27, 23, and 34 for CC-3pN, CC-5pN, CC-7pN, CC-10pN, and the control (without CC sensors), respectively). *P*-values are determined by one-way ANOVA with Tukey’s multiple comparison tests (\*\*\**p*<0.0001; ns>0.05: not significant).

Our novel coiled-coil (CC) force sensors are based on anti-parallel coiled-coils with short end-to-end distance that open in all or none fashion when they are under a force above a threshold pre-determined by their amino acid sequence (Fig. 1C) (Ren *et al*., 2025b). We have characterized sensors that open at 3, 5, 7, and 10 pN (called CC-3pN, CC-5pN, CC-7pN, and CC-10pN, respectively) (Table S1). Opening can be detected by exposure of reactive residues in the loop linking both α-helices. Here, we used the split-GFP readout system which uses the 11^th^ *β*-strand as the loop between the α-helices and the non-fluorescent GFP1-10 is expressed in the cytoplasm. When force opens the sensor, GFP11 becomes accessible, binds to GFP1-10, which becomes fluorescent. Since the GFP11/GFP1-10 heterodimer has a slow off-rate constant, transient opening of the coiled-coil produces a long-lasting signal. Therefore, it is used as a force recorder, which reports the maximum force that has been applied on the protein.

## Material and Methods

### Yeast strains and cell culture

*Schizosaccharomyces pombe* strains used in this study (Table S1) were maintained in YE5S (yeast extract supplemented with adenine, histidine, leucine, lysine, and uracil at 0.225 g/L). The strains were constructed by using CRISPR-Cas9 system (Ren *et al*., 2025a), and confirmed by sequencing of colony PCR with appropriate primer pairs.

### Growth assay

Fission yeast cells were cultured in YE5S until an optical density OD_595_∼0.3 and then resuspended to OD_595_=0.1 by water. The cells were placed on YE5S plates (20% agarose) with 10^1^, 10^2^, 10^3^ and 10^4^ dilutions. The plates were incubated for 48 hours at 32℃ or 37℃ incubator.

### Microscopy

Fission yeast cells grown to optical density OD_595_=0.3-0.6 were washed and resuspended in EMM5S (Edinburgh minimal medium supplemented with adenine, histidine, leucine, lysine, and uracil at 0.225 g/L. To acquire snapshots of cells using the split-GFP readout, resuspended cells were placed on 25% (w/v) gelatin pad. Fluorescence images were acquired every 0.5 µm for a total of 10 µm height (21 optical z-sections), using a spinning disk confocal microscope (Nikon, Japan) equipped with a 100x/NA1.45 objective, and 488 and 561 nm excitation lasers used to image mEGFP- and mScarlet-I-tagged proteins, respectively.

For time lapse imaging, resuspended cells were placed on a lectin-treated coverslip-bottomed dish. We used a spinning disk confocal microscope equipped with a 100x/NA1.45 objective (BC43, Oxford Instruments, UK). Z-stacks were acquired every 0.5 µm over 10 µm height (i.e., 21 optical sections), waiting 3-12 min interval between z-stacks, and using 488 and 561 nm excitation lasers.

### Quantification of force measurements with split-GFP readout

Because split-GFP is not completely irreversible, intensities can be used as a rough proxy of the fraction of sensors that spend some time open. In this study, we measured a force index as

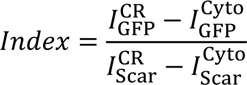

where *I* represents an average intensity in a contractile ring (I^CR^) or cytoplasm (I^Cyto^), and the subscripts represent GFP or mScarlet-I (Scar), using the maximum intensity projection of the acquired z-stacks. The index was analyzed using ImageJ (Schindelin *et al*., 2012; Schneider *et al*., 2012) and MATLAB (MathWorks).

### Analysis of ring constriction rates

We analyzed intensities from the maximum intensity projection of the acquired z-stacks. Ring diameters during fission yeast cytokinesis were obtained by tracking the fluorescently tagged ring marker Rlc1. Time courses of the ring diameters were fitted by a linear curve with the least-square method (MATLAB, MathWorks).

### Statistical analysis

Results are displayed as mean ± standard deviation (SD) unless otherwise noticed. P-values between two datasets were determined by Student’s *t*-test for two datasets, and by one-way ANOVA with Tukey’s multiple comparison tests for multiple groups using MATLAB (MathWorks), using the threshold p-values of 0.05 to determine statistical significance.

## Results

### Cdc12 transmits ∼3-6 pN of forces from its FH2 domain to the node through its N-terminus

To measure the force near the FH1 domain of Cdc12, we inserted coiled-coil (CC) force sensors that open under forces larger than 3, 5, 7, or 10 pN into A764 aa of fluorescently tagged Cdc12 (Fig. 1D,1E). Cdc12 harboring the CC-3pN and CC-5pN sensors at A764 emitted green fluorescence in the contractile rings, therefore reporting that the sensors are open and the force is larger than 3 and 5 pN, respectively. In contrast, CC-7pN, CC-10pN, and the control strain with no CC force sensors were not fluorescent in the GFP channel (Fig. 1E, 1F). The opened and closed force sensors in Cdc12 did not clearly affect cell growth (Fig. S1). To quantify the force measurements, the force index was obtained by Eq. (1). The force indexes of CC-3pN and CC-5pN were significantly larger than these of CC-7pN, CC-10pN, and the control strain (Fig. 1G). Accordingly, these results indicate that ∼6 pN of forces is transmitted at A764.

Next, we inserted the CC force sensors into several locations in N-terminus of Cdc12’s FH2, i.e. at T100 and A216 before the FH3 domain, L674 between the FH3 domain and FH1 domains, and K973 between the FH1 and FH2 domains (Fig. 2A). At T100 and A216, CC-3pN and CC-5pN were opened while CC-7pN and CC-10pN remained close (Fig. 2B-2E), indicating a maximum of ∼6 pN is transmitted through the N-terminal region of Cdc12. In contrast, at L674, all the force sensors tested (between 5 pN and 10 pN) remained closed, and CC-3pN emitted a weak GFP signal (Fig. 2F, 2G), demonstrating that the force at L674 is almost always very small but a very small fraction of Cdc12 experiences a force around 3 pN in rare occasions. At K973, CC-5pN resulted in significantly higher GFP signal than CC-7pN and CC-10pN (Fig. 2H, 2I). Altogether, Cdc12 transmits a force of ∼6 pN along its N-terminal end before the FH2 domain, except in a region right after the FH3 domain.

**Figure 2.**
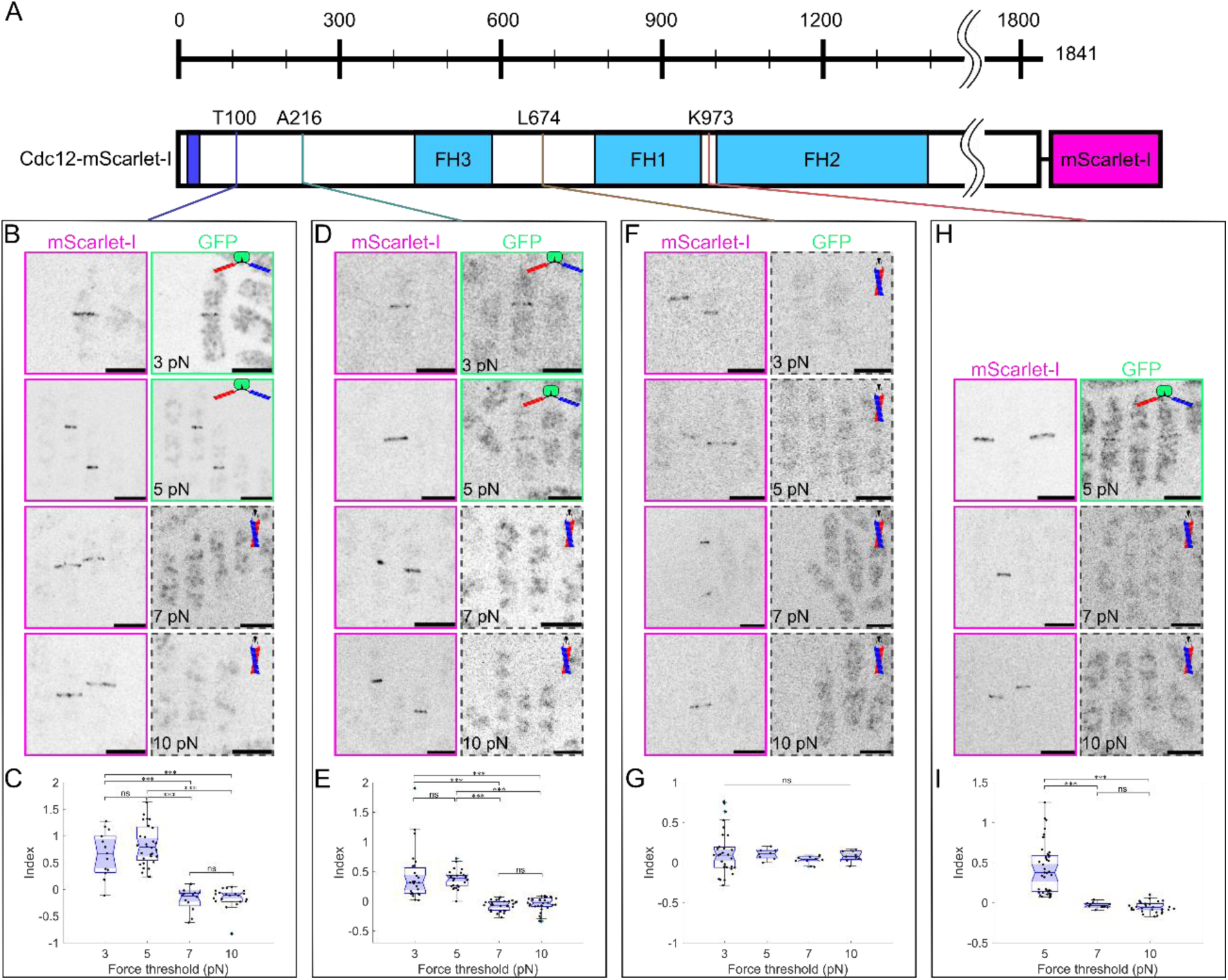
Forces on the upstream region of the FH2 domain of fission yeast Cdc12. **A** Schematics of the domain organization of the formin Cdc12 and insertion sites of the CC force sensors. **B, D, F, H** Fluorescence images of strains expressing fluorescently tagged Cdc12 (Cdc12-mScarlet-I) composed of the CC force sensors, and GFP1-10 replacing *pil1* gene. The coiled-coil force sensors with various force thresholds (3, 5, 7, or 10 pN) are respectively inserted at (**B**) T100, (**D**) A216, (**F**) L674, and (**H**) K973. Scale bars; 10 µm. **C, E, G, I** Quantification of the force measurements with the force index, represented as mean ± SD. **C** The force indexes at T100; *n*= 15, 32, 12, and 22 for CC-3pN, CC-5pN, CC-7pN, and CC-10pN, respectively. **E** Force indexes at A216; *n*= 27, 27, 28, and 30 for CC-3pN, CC-5pN, CC-7pN, and CC-10pN, respectively. **G** Force indexes at L674; *n*=30, 8, 10, and 9 for CC-3pN, CC-5pN, CC-7pN, and CC-10pN, respectively. **I** Force indexes at K973; *n*=42, 12, and 30 for CC-5pN, CC-7pN, and CC-10pN, respectively. *P*-values are determined by one-way ANOVA with Tukey’s multiple comparison tests (\*\*\**p*<0.0001; ns *p*>0.05: not significant) for (**C**), (**E**), (**G**), and (**I**).

### Force transmission in the N-terminal region of FH2 depends on Cdc12 binding to Cdc15 but not on Cdc12’s C-terminal region

Truncation of the first 503 aa of Cdc12, namely Δ 503-Cdc12, decreases the constriction rate of the cytokinetic ring (Coffman *et al*., 2013). Considering the slow constriction rate and the domain organization of Cdc12, we hypothesized that the force on the Δ503-Cdc12 formin mutant would decrease. We inserted the CC-5pN sensor with the split-GFP readout at A764 or K973 in Δ503-Cdc12. In both cases, the CC force sensors remained closed with no GFP signal (Fig. 3A, Fig. S2A), and their force indexes were significantly smaller than the indexes of similar sensors in the full-length Cdc12 (Fig. 3B, 2F and 2H). This result indicated that the first 503 aa of Cdc12 are necessary for the force transmission through the N-terminal end to the FH2 actin-binding domains.

**Figure 3.**
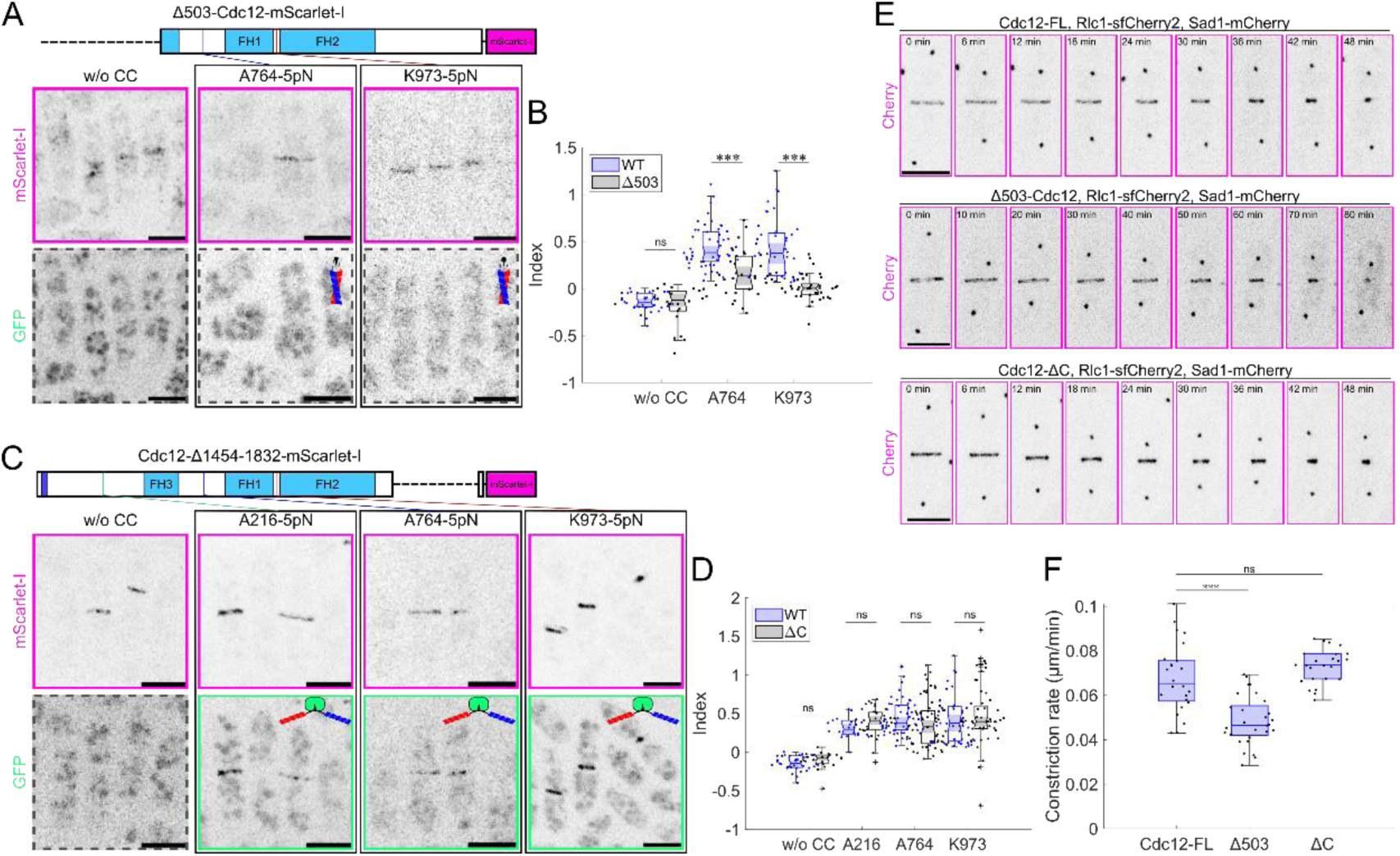
Forces in the upstream region of the FH2 domain of the formin Cdc12 depend on the N-terminal truncation. **A** Fluorescence images of the N-terminal 503 aa truncation mutants expressing the fluorescently tagged Cdc12 with and without the 5 pN coiled-coil force sensors (CC) at A764 and K973. Scale bars; 10 µm. **B** Quantification of the force measurements with the force index, represented as mean ± SD (*n*=33, 52, and 42 for the strains without CC, with CC-5pN at A764, and K973 in the full-length Cdc12 (blue), respectively; *n*=22, 30, and 37 for the strains without CC, with CC-5pN at A764, and at K973 in the Δ503-Cdc12 mutants (black), respectively). **C** Fluorescence images of the C-terminal tail (residues 1454-1832) truncation mutants of fluorescently tagged Cdc12 with and without 5 pN CC force sensors at A216, A764 and K973. Scale bars; 10 µm. **D** Quantification of the force measurements with the force index, represented as mean ± SD (*n*=33, 18, 52, and 42 for the strains without CC, with CC-5pN at A216, A764, K973 in the full-length Cdc12 (blue), respectively; *n*=24, 40, 64, and 52 for the strains without CC, with CC-5pN at A216, A764, K973 in the Cdc12-ΔC mutants (black), respectively). **E** Time courses of cytokinesis exhibited by the fluorescently tagged ring marker Rlc1 (Rlc1-sfCherry2) and the spindle pole body marker Sad1 (Sad1-mCherry) in the strains expressing the full-length Cdc12, Δ503-Cdc12, or Cdc12-ΔC. Scale bars; 10 µm. **F** Ring constriction rates determined by the time course of ring diameter in the strains expressing the full-length Cdc12, Δ503-Cdc12, or Cdc12-ΔC. *P*-values are determined by Student *t*-tests for (**B, D**) and by one-way ANOVA with Tukey’s multiple comparison test for (**F**) (\*\*\**p*<0.0001; ns *p*>0.05: not significant).

Next, we determined the role of Cdc12’s C-terminal force transmission to the node by measuring Cdc12’s N-terminal forces in a strains where most of Cdc12’s C-terminal region has been deleted, hereafter called Cdc12-ΔC. This formin mutant was previously shown to localize at rings and interphase spots (Yonetani *et al*., 2008). In contrast to Δ503-Cdc12, in the Cdc12-ΔC background, the force levels on Cdc12’s N-terminal region, A216, A764, and K973 (Fig. 3C, Fig. S2B), were similar as when the full length Cdc12 was expressed (Fig. 3D). Altogether, the force transmission in the C-terminal region of FH2 domain requires Cdc15 binding region in N-terminal of Cdc12 but is independent of Cdc12’s C-terminal region.

Next, we characterized the effects of the truncation on the constriction rates of the cytokinetic ring, in strains expressing the myosin-II regulatory light chain Rlc1 as a ring marker, and Sad1 as a spindle pole body marker, tagged with sfCherry2 and mCherry, respectively (Fig. 3E). We measured the ring constriction rate by fitting a linear function to the temporal evolution of the ring diameter (Fig. S3). The ring constriction rate of the Δ503-Cdc12 strain was approximately 30% slower than that of the full-length Cdc12. In contrast, the ring constriction rate in the Cdc12-ΔC strain was comparable with that of wild-type (Fig. 3F). These results support the idea that the N-terminus of Cdc12 is required for force transmission, but not its C-terminus.

### Force transmission on the C-terminus of Cdc12 is independent of Cdc12’s N-terminus and requires the C-terminal tail

The Cdc12’s C-terminal region after its FH2 actin-binding domains is predicted to be disordered. Since we showed the force transmission between the Cdc12’s N-terminus and the FH2 domain is independent of Cdc12’s C-terminal region (Fig. 3E,3F), we asked whether the C-terminal region of Cdc12 is under force. To test this hypothesis, we inserted our force sensors along the C-terminus at positions N1409, T1437, or G1576 (Fig. 4A). In all cases, CC-3pN and −5pN opened but CC-7pN and −10pN remained closed (Fig. 4B-4G). The truncation of Cdc12’s first 503 aa did not alter the force levels in the C-terminal region of Cdc12 (Fig. 4H, Fig. S4). These results suggest that force transmission to the C-terminal region of Cdc12 is independent of its N-terminal region.

**Figure 4.**
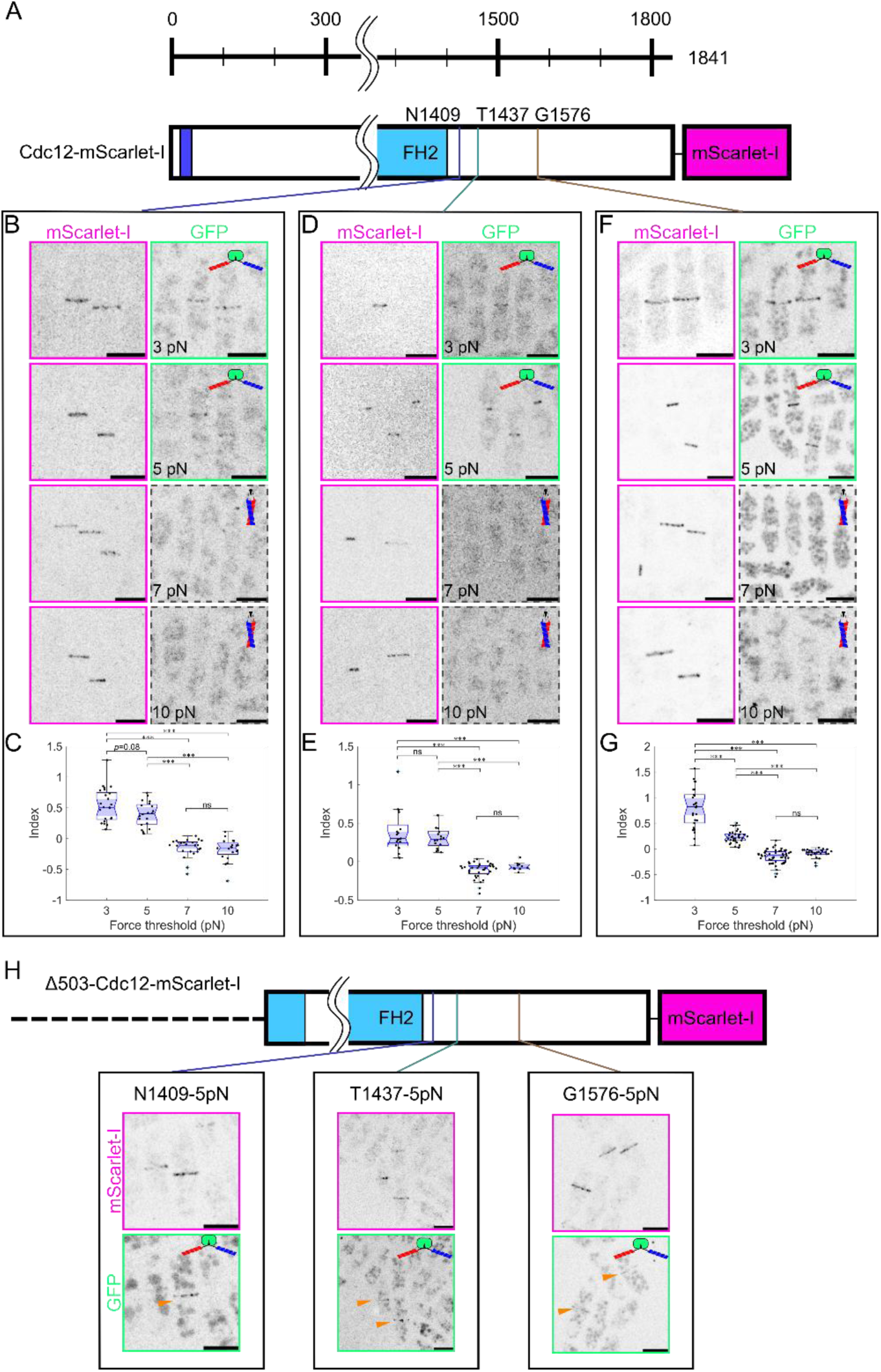
Force measurements with the coiled-coil (CC) force sensors along the C-terminal region of the FH2 domain of the formin Cdc12. **A** Schematics of the downstream region of the FH2 domain. **B, D, F** Fluorescence images of the strains expressing fluorescently tagged Cdc12 (Cdc12-mScarlet-I) composed of the CC force sensors, and GFP1-10 replacing *pil1* gene. The coiled-coil force sensors with various force thresholds (3, 5, 7, or 10 pN) are respectively inserted at (**B**) N1409, (**D**) T1437, and (**F**) G1576. Scale bars; 10 µm. **C, E, G** Quantification of the force measurements with the force index, represented as mean ± SD. **C** The force indexes at N1409; *n*= 25, 20, 23, and 20 for CC-3pN, CC-5pN, CC-7pN, and CC-10pN, respectively. **E** The force indexes at T1437; *n*= 17, 17, 35, and 10 for CC-3pN, CC-5pN, CC-7pN, and CC-10pN, respectively. **G** The force indexes at G1576; *n*=26, 33, 52, and 28 for CC-3pN, CC-5pN, CC-7pN, and CC-10pN, respectively. *P*-values are determined by one-way ANOVA with Tukey’s multiple comparison tests (\*\*\**p*<0.001; ns *p*>0.05: not significant). **H** Fluorescence images of the Δ 503-Cdc12 mutants expressing fluorescently tagged Cdc12 composed of the CC force sensors, and GFP1-10 replacing *pil1* gene. The 5pN CC force sensors are inserted at N1409, T1437, and G1576. Scale bars; 10 µm.

Next, we asked how the C-terminal region of Cdc12 is pulled with its N-terminal region-independent manner. To identify putative functional domains in the C-terminal region of Cdc12, we truncated part of the Cdc12 C-terminal region (residues 1423-1841), Cdc12-Δ1423-1841. On this background strain, we expressed high levels of this deleted C-terminal region (1423-1841) by fusing the sequence tagged with mScarlet-I to the end of highly expressed Hxk2 and separated the two sequences with the self-cleaving ERBV-1 2A peptide (Ren *et al*., 2023) (Fig. 5A). Overexpression of residues 1423-1841 clustered in the cytoplasm and localized to the contractile ring during cytokinesis (Fig. 5B). We interpreted this localization as a result of dimerization of the peptide with the oligomerization domain of Cdc12 (residues 1451-1538, (Bohnert *et al*., 2013)), known as the arginine- and serine-rich (RS) domain (Boucher *et al*., 2001). Indeed, the similar fragment lacking the RS domain did not form clusters (Fig. S5A, S5B). Overexpression of residues 1539-1841 also localized to the contractile ring during cytokinesis and aggregated in cytoplasm during interphase (Fig. 5C, 5D). This fragment also contains arginine and serine repeats, and deletion of the repeats led to clusters smaller than in the overexpression of the 1423-1841 fragment composed of the full RS repeats. In contrast, overexpression of residues 1577-1841 localized to the contractile ring and to filamentous structures that resemble actin bundles in the cytoplasm (Fig. 5E, 5F). Force measurements at G1576 in both the full-length Cdc12 and the overexpressed fragment (residues 1423-1841) resulted in ∼6pN of force (Fig. 4F, 4H, S5C-S5E). The replacement of residues 1577-1841 of Cdc12 with the maltose-binding protein (MBP) made the force at G1576 vanish (Fig. 5G, 5H, S5B). Altogether, our data show that residues 1577-1841 are necessary for force transmission from the C-terminal region of Cdc12 to the FH2 domain.

**Figure 5.**
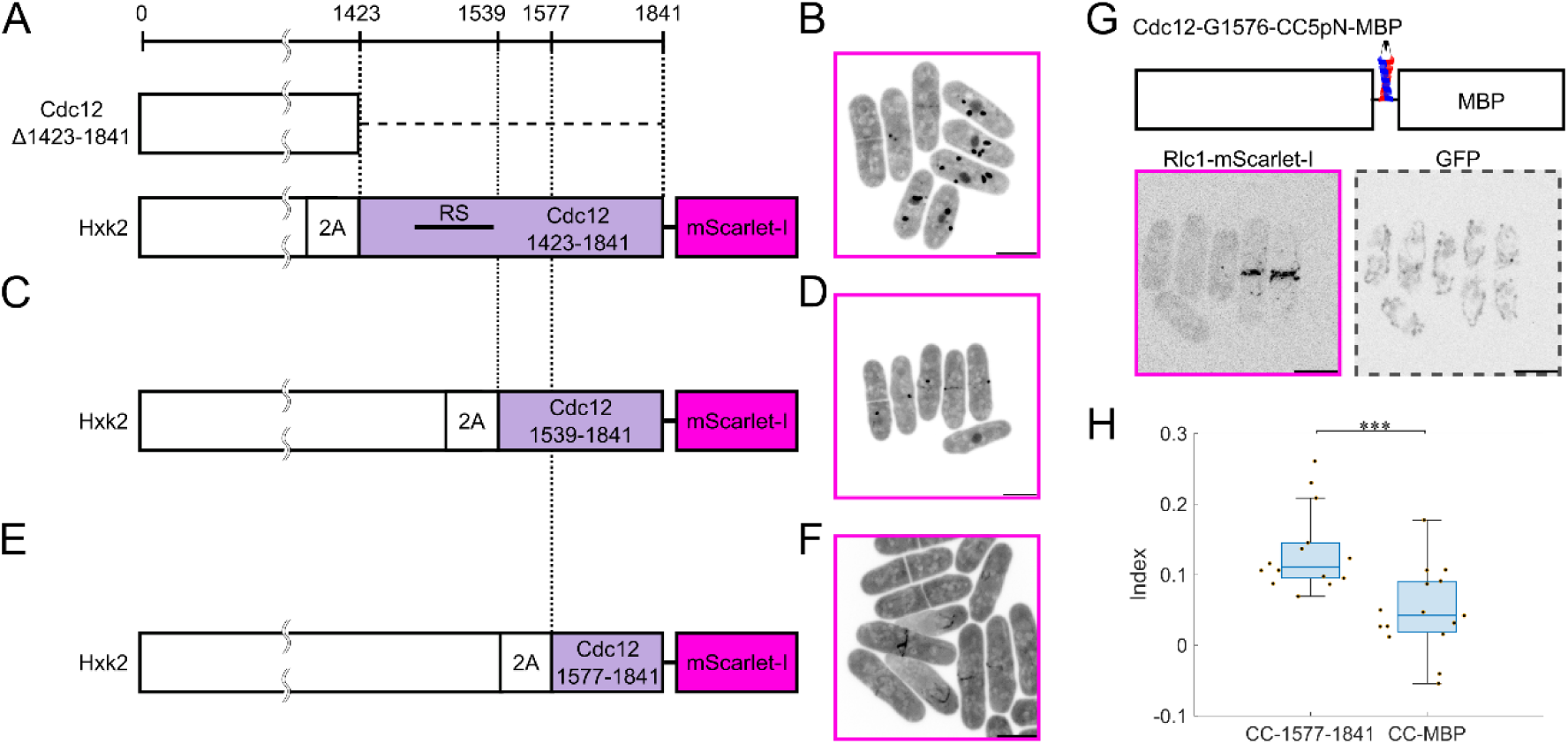
Distinct mechanisms of force transmission along the C-terminal region of the Cdc12 FH2 domain. **A, C, E** Schematics of highly expressed Hxk2 residues tagged with the self-cleaving ERBV-1 2A peptide and the Cdc12 C-terminal fragments; **A** Cdc12-1423-1841 containing the oligomerization domain (RS domain), **C** Cdc12-1539-1841, and **E** Cdc12-1577-1841, in the Cdc12-Δ1423-1841 strain. **B, D, F** Fluorescence images of the strains highly expressing the fragments of **B** Cdc12-1423-1841, **D** Cdc12-1539-1841, and **F** Cdc12-1577-1841. Scale bars; 10 µm. **G** Schematics of Cdc12 residues containing 5 pN CC force sensors at G1576 in the strain replacing the Cdc12-1577-1841 with the MBP. Fluorescence images of the strain expressing the fluorescently tagged ring marker Rlc1 (Rlc1-mScarlet-I), and GFP1-10 replacing *pil1* gene. Scale bars; 10 µm. **H** Quantification of the force measurements at G1576 in the wild-type Cdc12 and the mutant replacing the residues 1577-1841 with MBP. *P*-value is determined by Student’s *t*-test (\*\*\**p*<0.0001).

## Discussion

Actomyosin contractile rings play a central role in cell division. Although the molecular organization of contractile rings has been revealed (Laplante *et al*., 2016; Gerien and Wu, 2018; Pollard and O’Shaughnessy, 2019), the mechanisms of force transmission remained unclear. In this study, by using our coiled-coil force sensors (Ren *et al*., 2025b), we found that formin Cdc12 transmits forces of ∼6 pN along most of its length (Fig. 1, Fig. 2). In addition, our force measurements unveiled distinct and independent mechanisms for force transmission along the N- and C-termini of Cdc12. To transmit the forces from FH2 to the node, Cdc12’s N-terminus is required but not its C-terminus. Conversely, force transmission from Cdc12’s C-terminus to FH2 is not dependent on Cdc12’s N-terminus (Fig. 3A-3D).

The forces in the region between the N-terminus and FH2 domainwere ∼6 pN, except at L674 (Fig. 1E, 1G, 2B-2I). Whole ring tensions of ∼390 pN during interphase (Stachowiak *et al*., 2014) and ∼640 pN during ring constriction (Mcdargh *et al*., 2021) have been measured in fission yeast protoplasts. Since a cross-section of a contractile ring contains ∼50 actin filaments (Wu and Pollard, 2005; Courtemanche *et al*., 2016; Laplante *et al*., 2016; McDonald *et al*., 2017), we expect an average of ∼8-12 pN force per filament from these ring tension measurements. Moreover, simulations reported an average of ∼13 pN tensile force per filament (O’Shaughnessy and Thiyagarajan, 2018; Thiyagarajan *et al*., 2021). These experimentally and computationally estimated forces would transmit ∼4-6.5 pN on each subunit of formin Cdc12 homodimers, which is consistent with our force measurements of ∼6 pN. Our results are also in the same range as forces applied to nodes during their alignment into a contractile ring (∼6 pN), as estimated from first principles by analyzing the node motions (Vavylonis *et al*., 2008).

Formins’ mechano-sensitivity via FH1 and FH2 domains is a key feature of formin-mediated actin elongation. *In vitro* experiments revealed that piconewton forces accelerate actin polymerization of the human formin mDia1 (Jégou *et al*., 2013; Yu *et al*., 2017, 2018) and the budding yeast formin Bni1 (Courtemanche *et al*., 2013), possibly by helping the FH2 domain to switch from closed to opened conformation. In contrast, even sub-piconewton forces on fission yeast Cdc12 slow the actin polymerization *in vitro* (Zimmermann *et al*., 2017). The forces we measured between the FH1 and FH2 domains at K976 (∼6 pN, Fig. 2H) should be large enough to slow the force dependent actin elongation according to the *in vitro* observations (Zimmermann *et al*., 2017). Considering that the entropic spring-like disordered FH1 domain delivers profilin-actin molecules to the FH2 domain, this mechano-regulated inhibition of Cdc12 may be caused by stretching of the FH1 domain. Interestingly, we tried to insert between the FH1 and FH2 domains at K976 the 3 pN CC sensors, whose length when opened is approximately 2-fold larger than that of the other sensors used in our study (Table S1) (Ren *et al*., 2025b). However, this strain could not be constructed, possibly because this insertion was lethal. This result may support the importance of the distance between the FH1 and FH2 domains in Cdc12’s kinetics as shown previously(Courtemanche and Pollard, 2012; Bryant *et al*., 2017).

We were surprised to measure a force at L674 smaller than at other positions. We point out that even though the 3 pN force sensors remained closed, the index data spread in a distribution that is wider than the other ones (Fig. 2G), suggesting that forces in this region may be fluctuating. The L674 sensor insertion site is located between the putative dimerization domain (residues 550-630) adjacent to the FH3 domain (residues 321-503) and a coiled coil (residues 680-714) (Yonetani *et al*., 2008). These domains are conserved across many formins (Breitsprecher and Goode, 2013; Valencia and Quinlan, 2021). Moreover, Δ841-Cdc12 results in slow ring maturation and ring constriction, and reduces its level in the whole cell and at the contractile ring (Coffman *et al*., 2013), whereas Δ503-Cdc12 decreases only the ring constriction rate (Fig. 3E, 3F)(Coffman *et al*., 2013), meaning that the residues 504-841 help the ring maturation and Cdc12’s ring association. Considering the reduction of forces at A764 and K976 in the Δ503-Cdc12 strain (Fig. 3A, 3B), the putative dimerization domain and the coiled coil residues at 504-841 may interact with other Cdc12 domains or other proteins in a way that residues around L674 are protected from force.

Our force measurements in the C-terminal region of Cdc12 resulted in ∼6 pN of forces (Fig. 4). The N-terminal truncation of Cdc12 did not change the forces on the C-terminus (Fig. 4H, S4). These results suggest that there is a unique pulling mechanism in the C-terminal tail of Cdc12. Formin Cdc12 has a C-terminal disordered region after FH2 domain. While many formins are autoinhibited through the N-terminal Diaphanous inhibitory domain (DID) and the C-terminal Diaphanous autoregulatory domain (DAD) (Breitsprecher and Goode, 2013), this does not seem to be the case for the DID and DAD domains of formin Cdc12 (Yonetani *et al*., 2008). Therefore, we wondered how the C-terminal tail of Cdc12 can be stretched. In addition to the DAD, the C-terminal tail of Cdc12 contains an oligomerization domain with serine and arginine repeats (RS domain, residues 1451-1538) which contributes to the formation of linear actin bundles with weak actin-binding (Bohnert *et al*., 2013). This oligomerization may contribute to the transmission of pulling forces through the region downstream of the FH2 domain (Fig. 4A-4E). However, our force measurements at G1576, in C-terminal to the RS domain, also resulted in ∼ 6 pN of forces (Fig. 4F, 4G). In addition, the Cdc12’s C-terminal tail itself opened the 5 pN force sensors at G1576, showing a similar phenotype to the previous report (Yonetani *et al*., 2008), but more dramatic aggregations upon the overexpression (Fig. S5C). These results imply the existence of one or multiple other binding motifs different from the RS repeats in the region downstream of G1576. Indeed, the overexpression of the residues 1577-1841 localized to the contractile rings and structures that resemble actin cables (Fig. 5E, 5F). By replacing residues 1577-1841 with MBP, the force sensors at G1576 remained closed (Fig. 5G, 5H). These results support the idea that the residues 1577-1841 associate with the contractile rings, contributing the stretch of the C-terminal region of Cdc12.

## Acknowledgements

This study was partly supported by NIH grants R21GM132661, R01GM115636 and the Research Corporation for Scientific Advancement grant SA-CMC-2021-037 and JSPS (Japan Society for the Promotion of Science) 22J00060. We thank the Yale West Campus Imaging Core for resources for microscopy and Keck DNA Sequencing Facility at Yale for helping DNA confirmation of strains.

## Author contributions

Takumi Saito: Writing – original draft, Visualization, Validation, Methodology, Funding acquisition, Formal analysis, Data curation. Yuan Ren: Writing – review & editing, Visualization, Validation, Methodology, Data curation, Conceptualization. Julien Berro: Writing – review & editing, Validation, Supervision, Software, Resources, Project administration, Investigation, Funding acquisition, Conceptualization.

## Competing interests

Julien Berro and Yuan Ren have a pending patent application PCT/US2023/069505. Takumi Saito declares no competing financial interests or personal relationships that could have appeared to influence the work reported in this paper.

## Supplemental materials of

**Figure S1.**
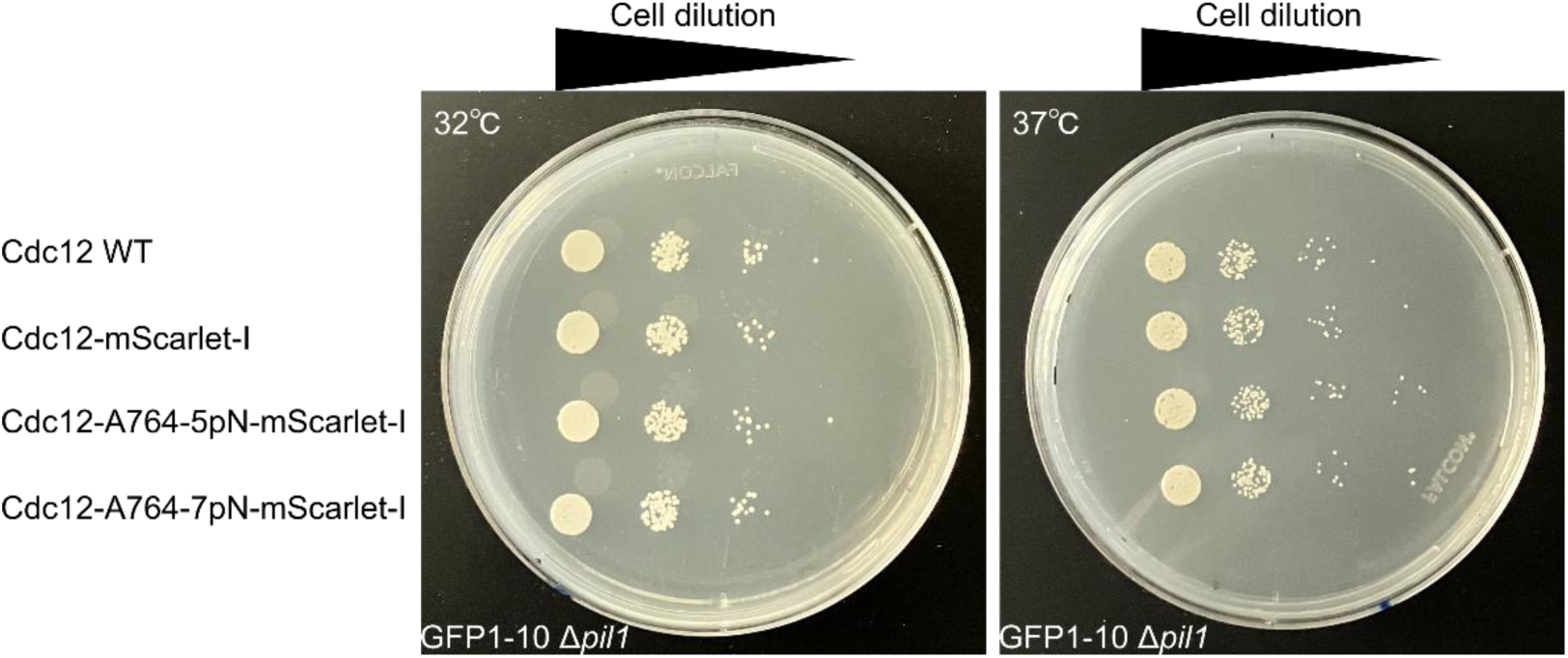
Growth assay of representative *S. pombe* strains. GFP1-10 is expressed from the *pil1* gene locus. The coiled-coil force sensors are inserted at A764 of fluorescently tagged Cdc12. The cells are spotted on YE5S plates after 1 to 10^1^, 10^2^, 10^3^ and 10^4^ dilutions and incubated for 48 hours at 32℃ or 37℃.

**Figure S2.**
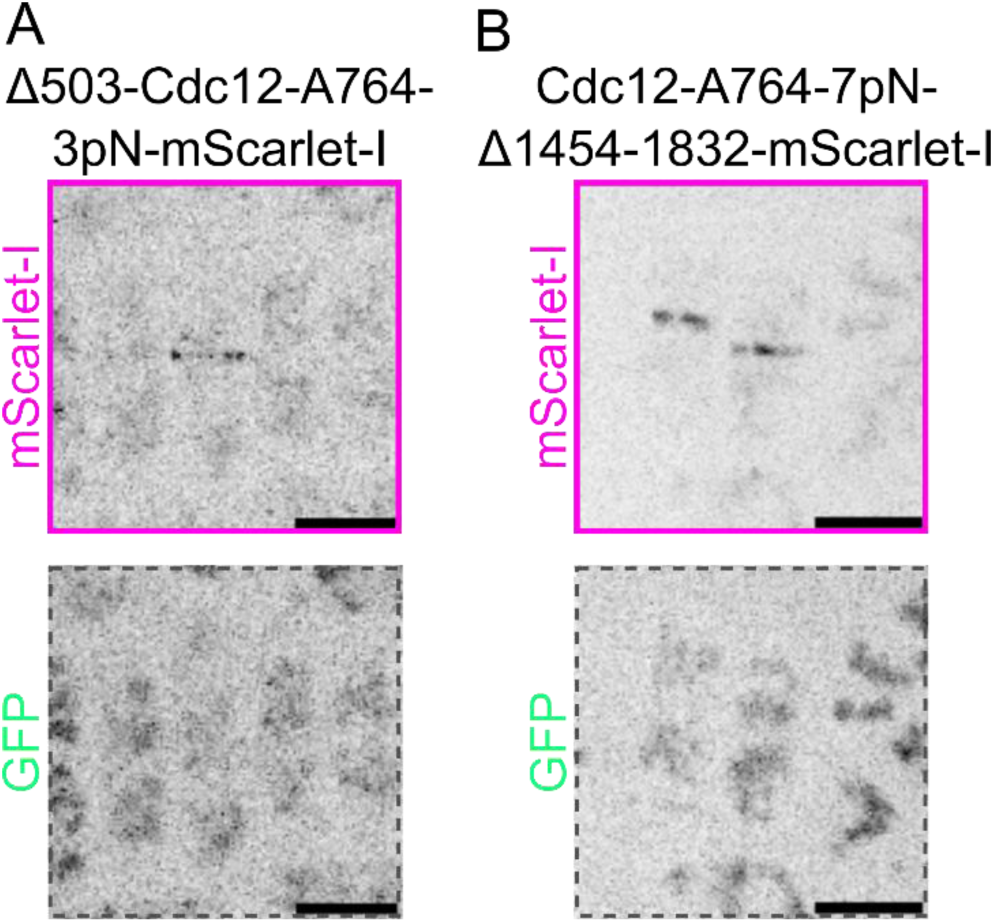
Force measurements at A764 in Δ503-Cdc12 and Cdc12-Δ1454-1832 strains with (**A)** the 3 pN and (B) the 7 pN coiled-coil force sensors, respectively. Scale bars; 10 µm.

**Figure S3.**
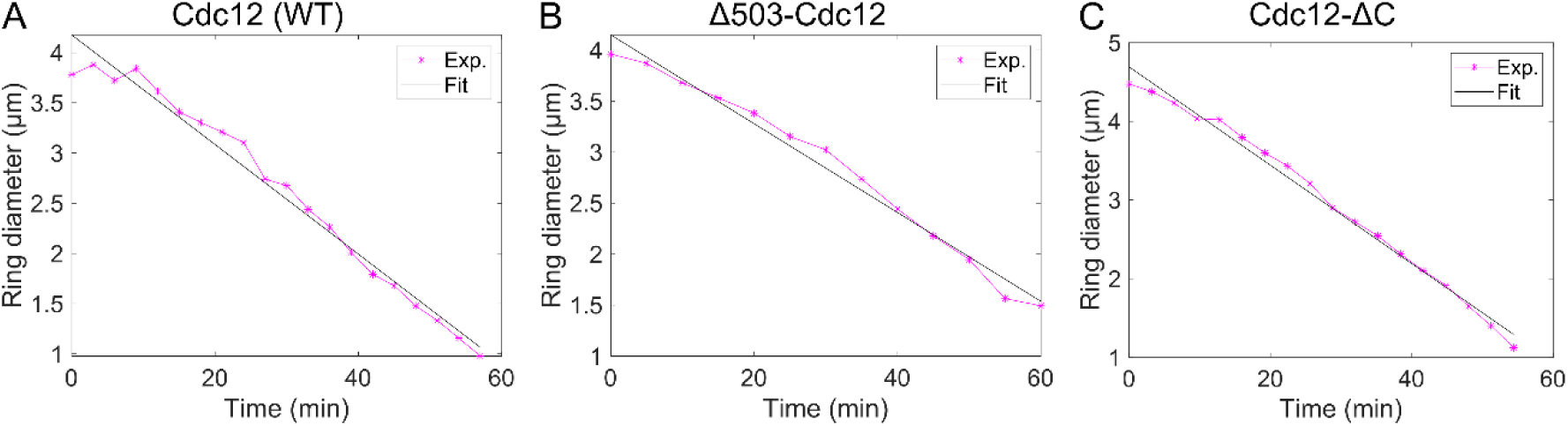
Representative plots of ring constriction of the strains expressing **A** the wild type Cdc12, **B** Δ503-Cdc12, or **C** Cdc12-ΔC (1454-1832 aa).

**Figure S4.**
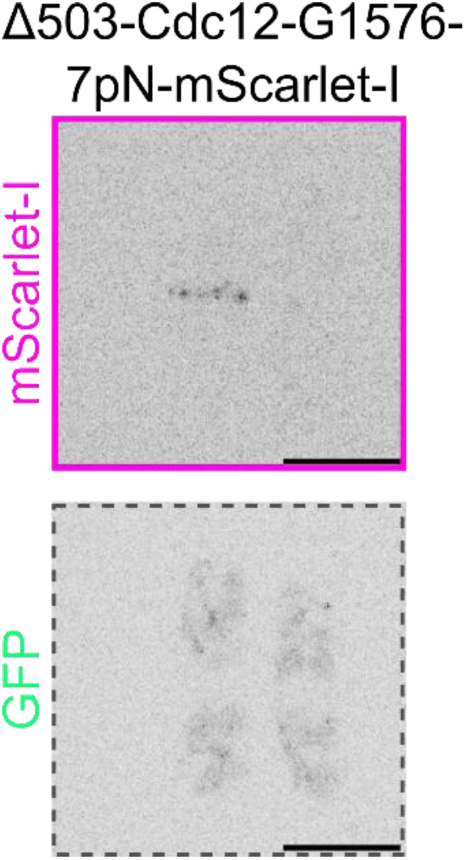
Snap shots of the strain expressing Δ503-Cdc12 with the 7 pN coiled-coil force sensors at G1576. Scale bars; 10 µm. 21

**Figure S5.**
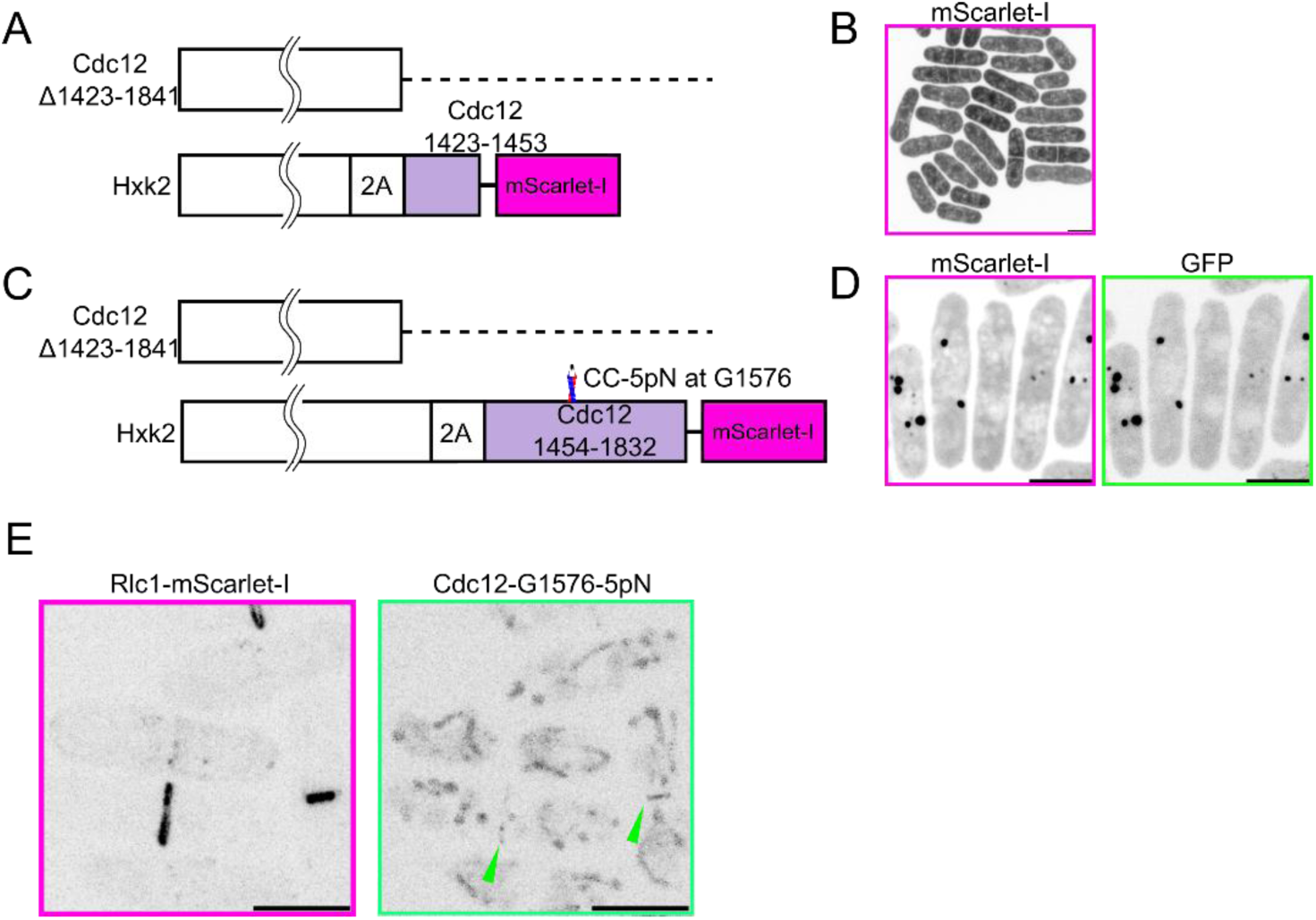
Overexpression of Cdc12 C-terminal regions. **A** Schematics of 1423-1841 aa truncation of Cdc12 and the Cdc12 C-terminal fragment (residues 1423-1453) which is expressed in combination with the highly abundant Hxk2 and separated from it via the self-cleaving ERBV-1 2A peptide. **B** Fluorescence image of the strain highly expressing the fragment of Cdc12-1423-1453 (A). Scale bar; 10 µm. **C** Schematics of 1423-1841 aa truncation of Cdc12 and the Cdc12 C-terminal fragment (residues 1454-1832) containing the 5 pN force sensors (CC-5pN) at Cdc12-G1576, expressed in combination with the highly expressed Hxk2 and separated from it with the self-cleaving ERBV-1 2A peptide **D** Fluorescence images of the strain highly expressing the fragment of Cdc12-1454-1832 with the CC-5pN at G1576. Scale bars; 10 µm. **E** Fluorescence images of the strain expressing mScarlet-I tagged Rlc1 and the full length Cdc12 with the 5 pN sensors at G1576. Scale bars; 10 µm. 36

**Table S1.**
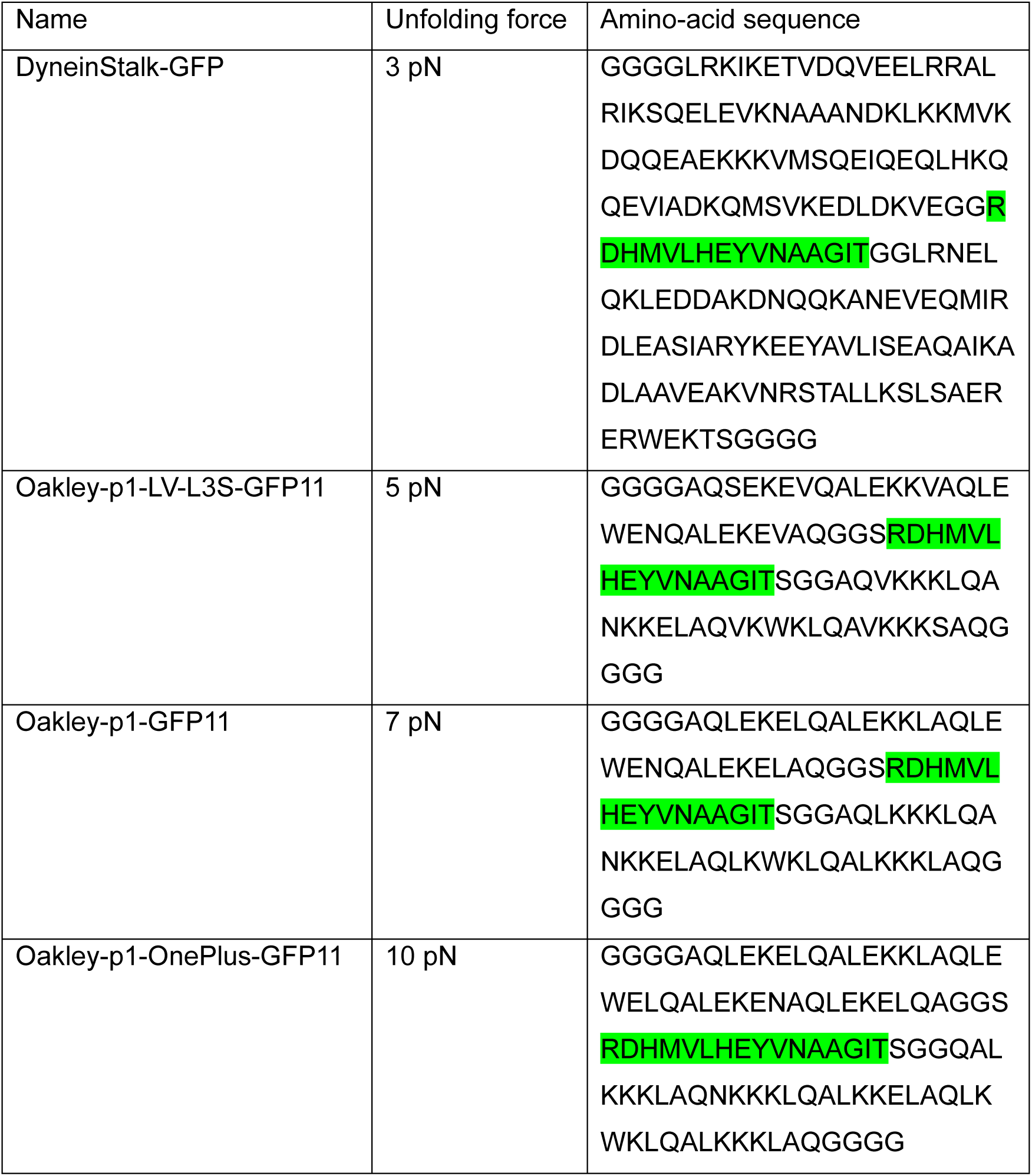
Coiled-coil force sensors with the GFP11 linker highlighted in green.

**Table S2.**
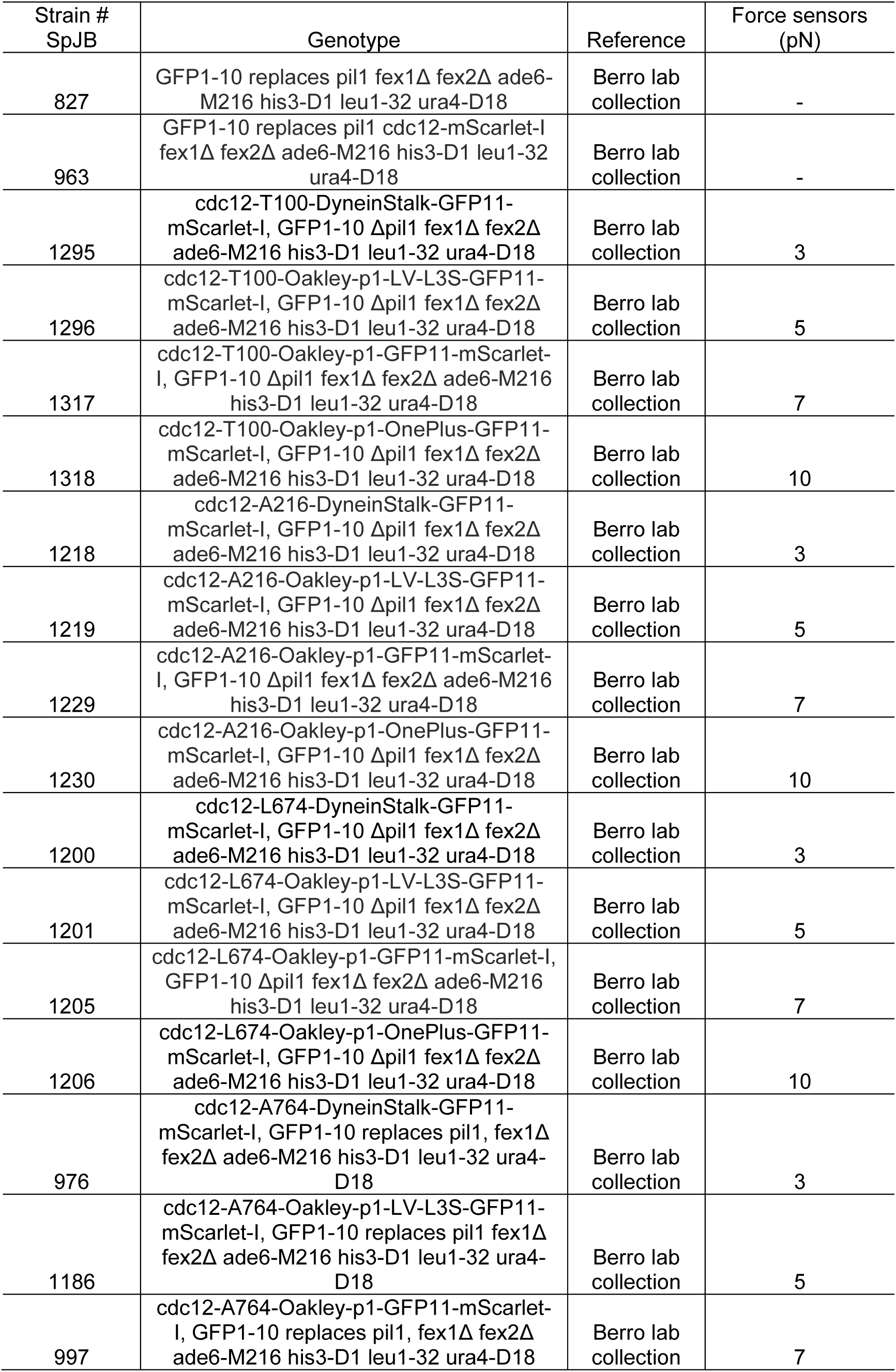
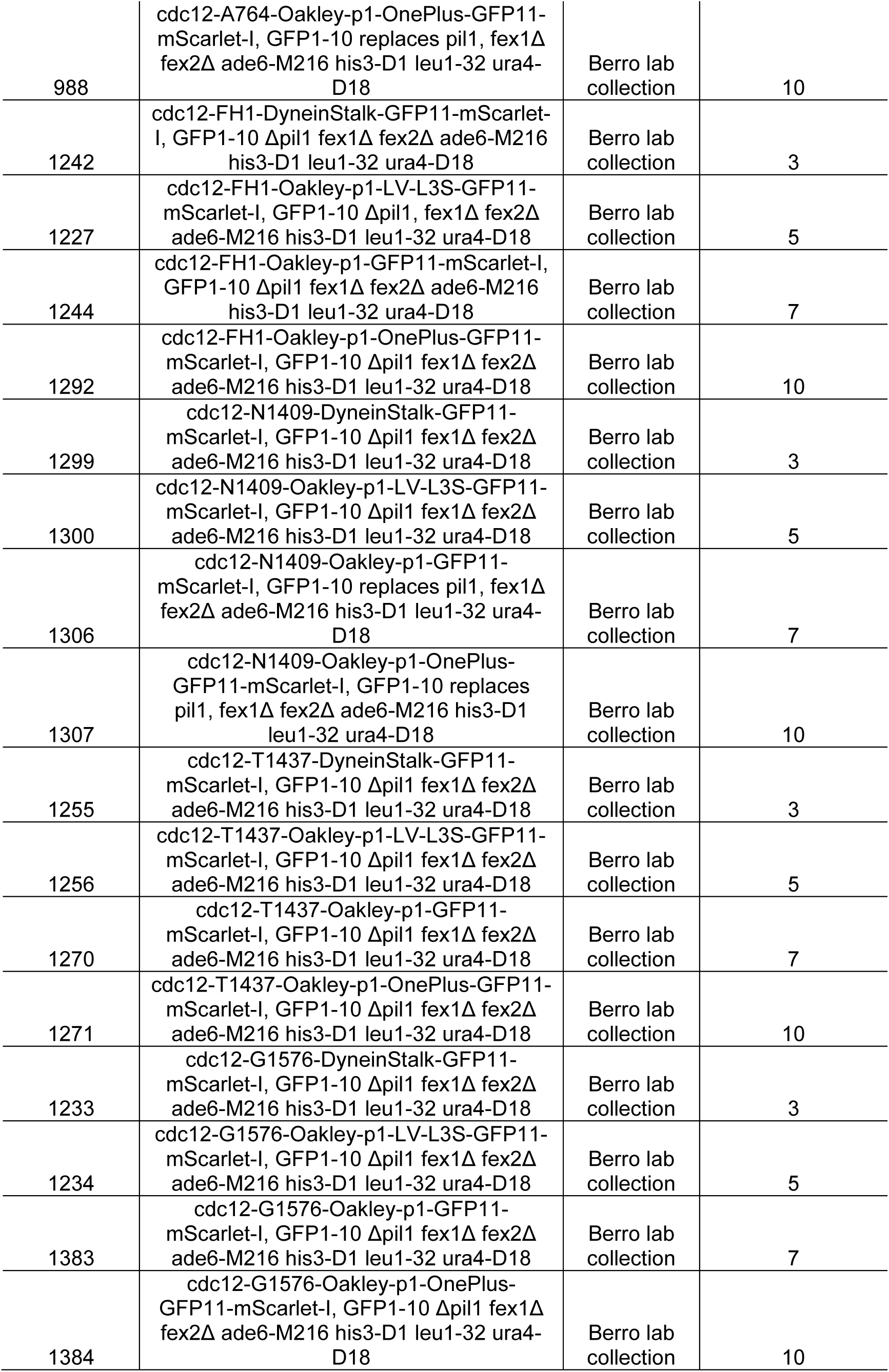
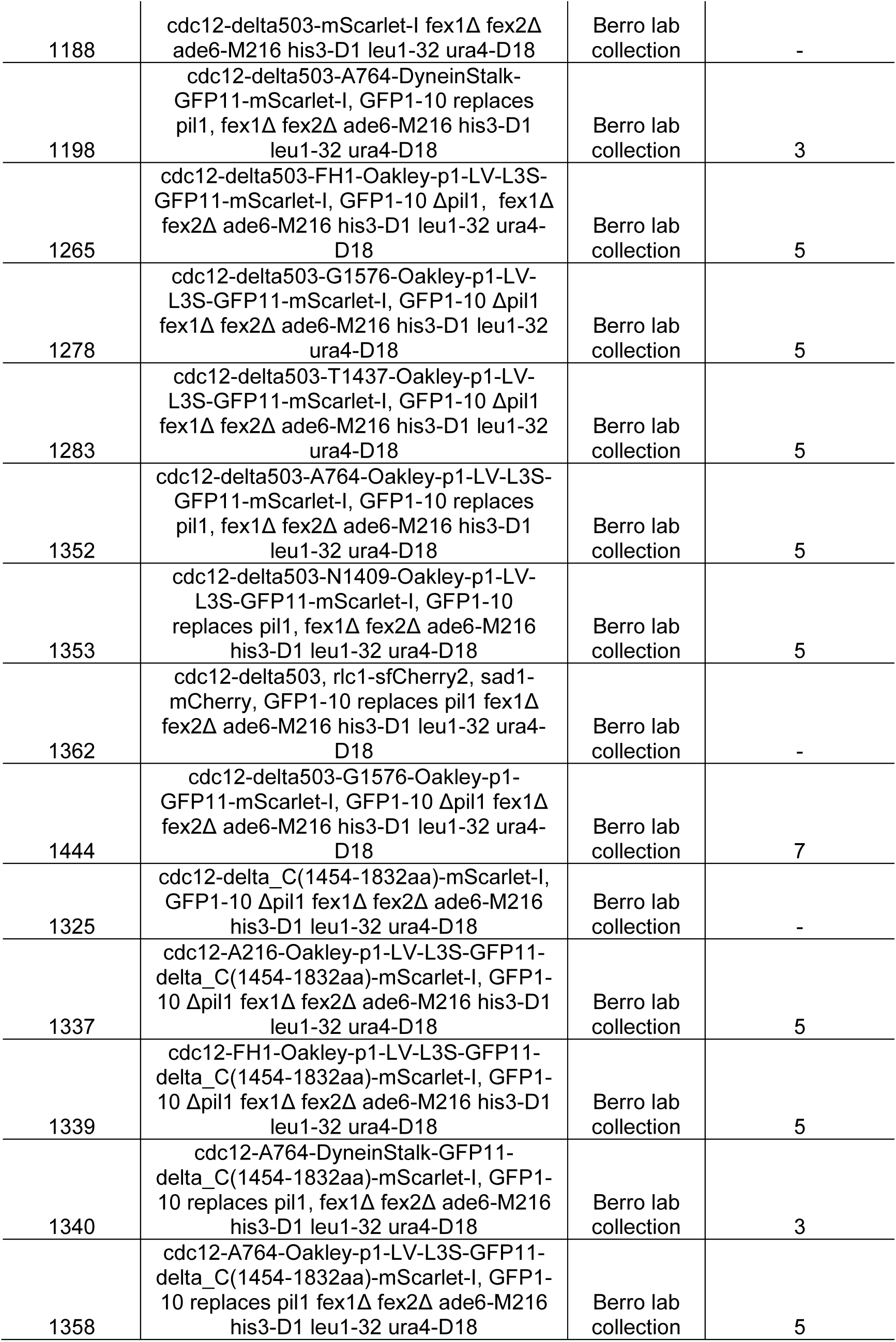
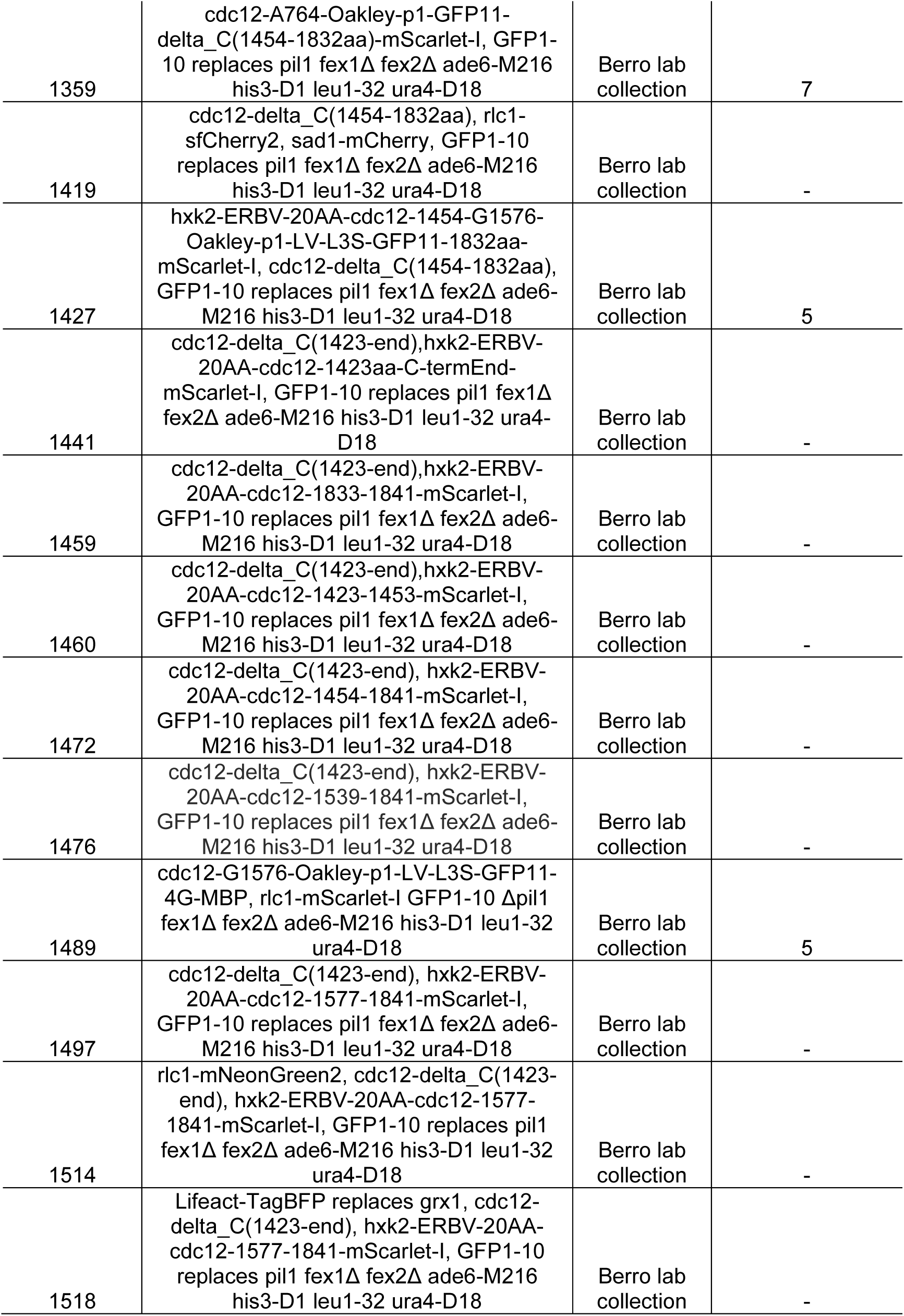
*S. pombe* strains used in this study.

